# Antifungal effects of PC945, a novel inhaled triazole, on *Candida albicans*-infected immunocompromised mice

**DOI:** 10.1101/2020.07.27.222661

**Authors:** Yuki Nishimoto, Kazuhiro Ito, Genki Kimura, Kirstie A. Lucas, Leah Daly, Pete Strong, Yasuo Kizawa

## Abstract

Although *Candida spp.* are frequently detected in fungal cultures of respiratory secretions, their presence is normally assumed to reflect benign colonization. However, there is growing evidence that *Candida spp.* are involved in the pathogenesis of respiratory diseases such as bronchiectasis and asthma. The aim of this study is to investigate the *in vitro* and *in vivo* effects of a novel antifungal triazole, PC945, optimised for topical delivery, against *C. albicans*. In temporarily neutropenic, immunocompromised mice, *C. albicans* (529L [ATCC^®^MYA4901™] strain), inoculated intranasally, induced acute lung injury and death, associated with higher fungal burden and cytokine induction in the lung. PC945 saline suspension, dosed intranasally once daily, starting one day post candida inoculation, dose-dependently (0.56 ~ 14 μg/mouse) improved survival rate and inhibited fungal load in the lung on Day 5 post inoculation. These effects by PC945 were 7 ~ 25-fold more potent than those of voriconazole, despite being of similar *in vitro* antifungal activity versus this strain. Furthermore, extended prophylaxis with low dose PC945 (0.56 μg/mouse) was found to inhibit fungal load more potently than the shorter treatment regimens, suggesting antifungal effects of PC945 accumulated on repeat dosing. In addition, antifungal susceptibility testing on 88 candida isolates *(C. albicans, C. parapsilosis, C. tropicalis, C. lucitaniae, C. glabrata, C. guilliermondii)* revealed that PC945 has potent effects on *Candida* species broadly. Thus, PC945 has the potential to be a novel topical therapy for the treatment of *C. albicans* pulmonary infection in humans.

## Introduction

*Candida spp.* are often detected from fungal cultures or mycobiome analysis of respiratory secretions, but are not usually treated since, even if detected persistently, the presence is assumed to be benign colonization unless invasive candidiasis in deeply immunocompromised subjects is suspected. Thus, although antifungal therapy is standard treatment for *Candida* infections observed in other organs such as the vagina and skin, no practical guidelines to treat *Candida* infection in the respiratory tract, except for invasive candidiasis, have been developed (1, 2).

However, there is growing evidence that *Candida spp*. may be causally involved in chronic respiratory diseases as well as invasive candidiasis via the respiratory tract. For example, *Candida albicans* as well as *Aspergillus fumigatus* are often detected in patients with non-cystic fibrosis bronchiectasis, who show compromised mucociliary clearance (3, 4). Candida can also cause pneumonia with a severe clinical course (5, 6). Johnson reported 11 cases of chronic candida bronchitis patients who had persistent sputum production despite antibiotic treatment, where antifungal therapy led to a good or excellent clinical response (improved respiratory symptoms and sputum production) within 3 weeks (2). Corticosteroid treatment is known to facilitate *C. albicans* colonization of the respiratory tract (7) as well as oral/oesophageal candidiasis (8). Candida itself also causes allergic bronchial pulmonary mycosis (9) and sensitisation to *C. albicans*, while not an aeroallergen, is seen in up to 10% of individuals with mild stable asthma and 33% of patients with severe asthma (10). As well as worsened sputum production and bronchitis, persistent *C. albicans* was associated with worse post-bronchodilator forced expiratory volume (FEV1), more frequent bronchiectasis and more hospital-treated exacerbations in non-cystic fibrosis patients with bronchiectasis (11). Furthermore, candida airway colonization has been shown to be associated with prolonged duration of mechanical ventilation and ICU/hospital length of stay. The overall hospital mortality in this group was significantly higher than patients with non-Candida albicans fungal infections (12, 13).

Beta-glucan, mannan and chitin, components of the yeast cell wall, may act as proinflammatory factors in the lung (14). In addition, Candida is known to interplay with both Gram-positive and Gram-negative bacteria in the environmental biofilm, through quorum-sensing molecules (15). *In vivo*, *Candida albicans* infection was found to worsen pneumonia caused by *Pseudomonas aeruginosa*, *Escherichia coli*, and *Staphylococcus aureus* in rats, enhancing the production of inflammatory cytokines in the lung, and inhibiting phagocytosis by alveolar macrophage (16–18).

Systemic triazole therapy is the basis of treating infections with pathogenic fungi but the adverse effects of itraconazole, voriconazole and posaconazole are well characterised and thought to be a consequence of their pharmacological effects in host tissues (19, 20). Furthermore, notable drug interactions for voriconazole due to the inhibition of hepatic P450 enzymes make clinical use challenging and indeed the variability in exposure of the triazoles via the oral route necessitates the need for close therapeutic drug monitoring and limits the use of triazole therapy prophylactically in at risk groups (21, 22). In addition, fungus colonization occurs in pre-existing lung cavities due to tuberculosis or on airway surfaces where fungi first deposit and grow, and is difficult to deliver and maintain appropriate local levels of anti-fungal agents after oral treatment (23). Furthermore, resistance to antimicrobial agents is an emerging problem worldwide. Although high levels of, and continuous exposure to, antimicrobial agents is known to decrease the risk of mutation induction leading to resistance (24), systemic treatment hardly achieves continuous high levels of exposure (for example, AUC/MIC, >60 (24)) in lung cavities in all patients although the level might exceed MIC values (23, 25). Thus, inhaled treatment has several potential advantages versus oral/systemic treatment which alter the risk benefit ratio of treatment favourably. Tolman and colleagues have demonstrated that prophylaxis with an aerosolized aqueous intravenous formulation of voriconazole significantly improved survival and limited the extent of invasive disease, as assessed by histopathology, in an invasive pulmonary murine model (26). In human studies, aerosolization of voriconazole showed beneficial effects but it required frequent and high doses as voriconazole was not optimised as an inhaled medicine (27, 28).

PC945, 4-[4-(4-{[(3R,5R)-5-(2,4-difluorophenyl)-5-(1H-1,2,4-triazol-1-ylmethyl)oxolan-3-yl]methoxy} −3-methylphenyl)piperazin-1-yl]-N-(4-fluorophenyl)benzamide, is a novel antifungal triazole designed to deliver; high local concentrations, retention in cells offering a long duration of action, minimal systemic exposure with poor oral availability and high protein plasma binding (29, 30). PC945 has been shown to have potent *in vitro* and *in vivo* antifungal activity on *A. fumigatus* isolates with inhibition of the enzyme lanosterol 14α-demethylase (CYP51A1) and also demonstrated that topical once daily treatment of PC945 was highly effective versus fungal burden in lung tissue and several biomarkers (galactomannan, cytokines) in serum and bronchoalveolar lavage fluid (BALF) in temporarily neutropenic mice infected with *A. fumigatus* when compared with intranasally treated posaconazole and voriconazole. Interestingly, the first case report describing the successful use of PC945 to treat a refractory *Aspergillus* bronchial anastomotic infection and tracheobronchitis in a lung transplant on top of standard care has recently been published (31). Earlier studies showed anti-fungal activities of PC945 against *C. auris* (32) and limited isolates of *Candida spp. (33)*.

Thus, the aim of this study is to evaluate in vitro antifungal effects of PC945 by broth microdilution assay with an extended collection of *Candida spp.* and investigate the *in vivo* antifungal effects of intranasally dosed PC945 in temporarily neutropenic mice infected with *C. albicans*.

## Materials and methods

### Antifungal agents

PC945 was synthesised by Sygnature Discovery Ltd (Nottingham, UK), while voriconazole (Tokyo Chemical Industry UK Ltd., Oxford, UK) and posaconazole (Apichem Chemical Technology Co., Ltd., Zhejiang, China) were procured from commercial sources. For *in vitro* antifungal assays, stock solutions of test agents were prepared in DMSO (2000 μg/ml). For *in vivo* studies test agents were suspended in physiological saline.

### *In vivo* antifungal activity against *C. albicans* infection

Specific pathogen-free A/J mice (male, 5 weeks old) were purchased from Sankyo Labs Service Co. Ltd. (Tokyo, Japan) and adapted for 1 week in a temperature (24 ± 1°C) and humidity (55 ± 5%) controlled room, under a 12-h day-night cycle. The mice were reared on a standard diet and tap water *ad libitum*. A/J mice were used for *A. fumigatus* infection and proved to be more efficiently infected as described previously (21). Animals were then dosed with hydrocortisone (Sigma H4881, 125 mg/kg, subcutaneously) on days 3, 2 and 1 before infection, and with cyclophosphamide (Sigma C0768; 250 mg/kg, intraperitoneally) two days before infection to induce temporary neutropenia as previously reported (22). Both hydrocortisone and cyclophosphamide were diluted with physiological saline. To avoid bacterial infection, drinking water was supplemented with tetracycline hydrochloride (Sigma T7660; 1 μg/ml) and ciprofloxacin (Fluka 17850; 64 μg/ml). *C. albicans* (ATCC MYA4901 [529L]), purchased from the American Type Culture Collection (Manassas, VA, USA), was grown on Sabouraud dextrose agar (Difco Laboratories Ltd., West Molesey, UK) plates for 3 days at 35°C. Fungal colonies were aseptically dislodged from the agar plates and suspended in physiological saline. On the day of infection, yeast counts were assessed by haemocytometer, and the inoculum was adjusted to obtain concentrations of 5 × 10^7^/mL of sterile physiological saline. On Day 0, the suspension was administered intranasally (total volume 50 μL/mouse; 25 μL per each nostril), so 2.5 × 10^6^ yeast was inoculated per mouse.

PC945 or voriconazole, suspended in physiological saline, were administered intranasally (total volume 35 μL/mouse; approximately 17.5 μL each to each nostril) once daily from Day 1 to Day 5 post inoculation as shown in Fig 1. To investigate extended prophylaxis, PC945 was administered intranasally once daily, on days −7 to 0 and the effects were compared with treatment on days −1 to 0. As the injection volume was fixed and body weight changed daily, especially after infection, the accurate dose unit was μg/mouse. However, we also calculated estimated dose as mg/kg, assuming the average body weight of 20g just before treatment and 60% exposure after intranasal dosing. Therefore, 35 μl injections of 0.016, 0.08, 0.4, 2, 10 mg/ml were equivalent to 0.56, 2.8, 14, 70, 350 μg/mouse, respectively, corresponding to approximately 0.028, 0.14, 0.7, 3.5 and 17.5 mg/kg, respectively (S Table 1).

**Fig 1.**
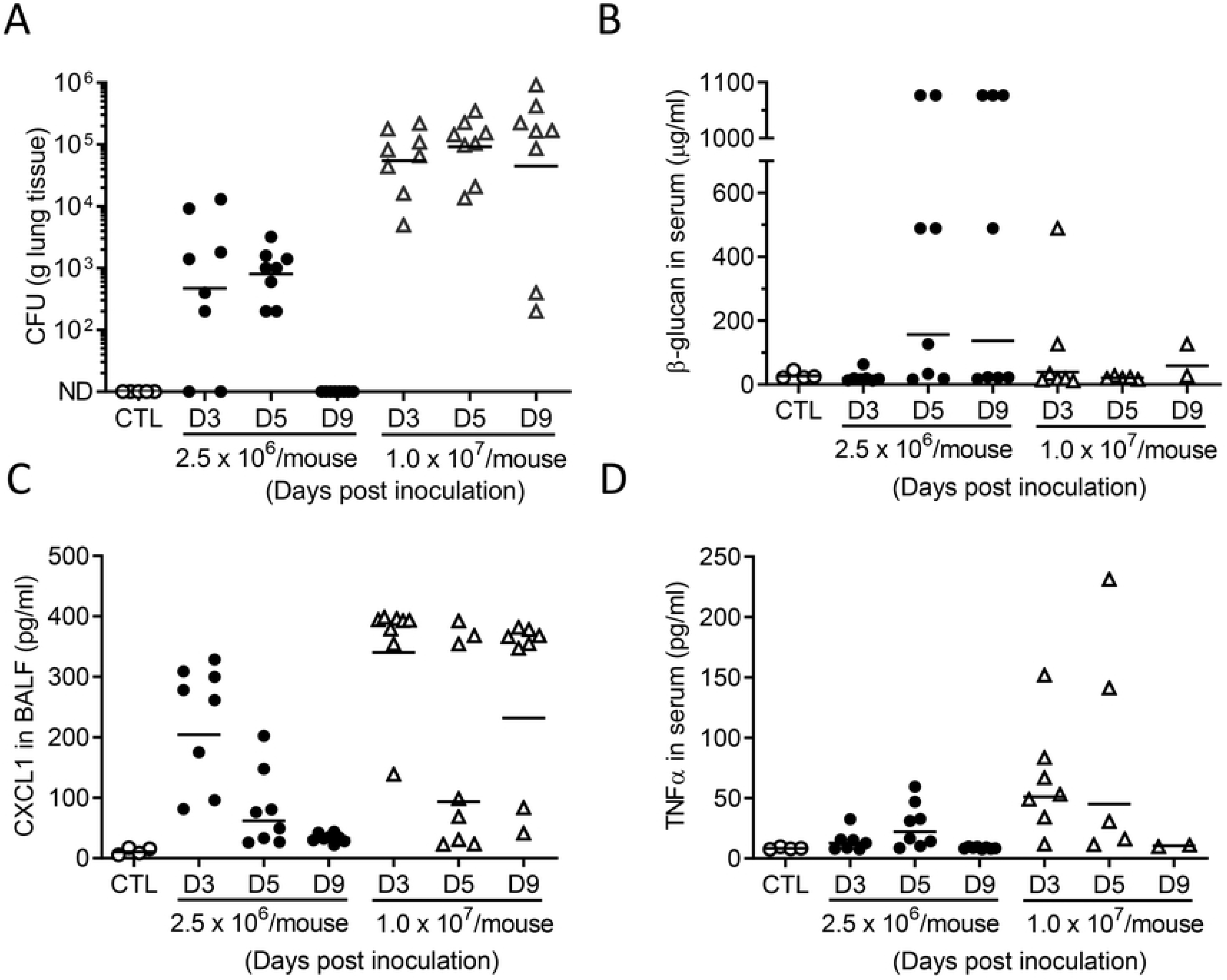
Experimental protocol with dosing regimen. *C. albicans* (529L) and compounds (aqueous suspension) were given intranasally.

Deaths and the body weights of surviving animals were monitored daily for 5 days. A body weight loss of > 20% or more, compared with an animal’s weight a day before, or a mouse death, were defined as “drop-out” events. Bronchoalveolar lavage fluid (BALF) was collected through cannulated tracheas using physiological saline (34) on day 5 post infection (6 hrs after the final treatment) or on the day that the mouse dropped out of the study. Blood was then collected via cardiac puncture except for dead mice, and left lung tissue was removed to prepare a tissue homogenate. All animal studies were approved by the Ethics Review Committee for Animal Experimentation of Nihon University. *C. albicans* studies were approved by the Microbial Safety Management Committee of Nihon University School of Pharmacy (E-H25-001).

### Biomarker analysis

For semi-quantitative tissue fungal load analysis, 100 mg of lung tissue (left lung) was removed aseptically and homogenized in 1.0 mL of sterile saline and kept on ice to limit fungal growth of in homogenates until all samples were processed. Homogenates were prepared using a mini cordless CG-4A homogenizer (Funakoshi Ltd., Tokyo, Japan) using mild conditions (2 repeated cycles of 10 seconds homogenization and 2 min resting on ice) before centrifugation at 1400 x g for 1 min (room temperature). The supernatants obtained were serially diluted using sterile physiological saline and plated on Sabouraud agar plates (50 μL/plate), and then incubated at 35±1°C for 72 to 96 hrs. The colonies of *C. albicans* on each plate were counted and corrected to the dilution factor, and the fungal titre is presented here as CFUs per gram of lung tissue. The levels of CXCL1 in BALF and TNFα in serum were determined using Quantikine^®^ mouse ELISA kits (R&D systems, Inc., Minneapolis, MN, USA). The *Candida* β-glucan concentration in BALF or serum was determined with Glucatell kits (Associates of Cape Cod, Inc., Massachusetts, USA). For histology, the right lung was fully inflated by intratracheal perfusion with 10% paraformaldehyde in PBS. Lungs were then dissected free and placed in fresh paraformaldehyde solution. Routine histological techniques were used to paraffin-embed the tissues, and 4 μm sections of whole lung were stained with either haematoxylin and eosin or Grocott stain (Silver stain kit, Sigma HT100A).

### *Candida* Strains and MIC determination

Antifungal susceptibility testing for 88 *candida* isolates *(C. albicans* [37]*, C. parapsilosis* [17]*, C. tropicalis* [7]*, C. lucitaniae* [11]*, C. glabrata* [10]*, C. guilliermondii* [6]) was performed in accordance with the guidelines in the Clinical and Laboratory Standards Institute (CLSI) M27-A4 document (4^th^ edition)(35) and the European Committee on Antimicrobial Susceptibility Testing (EUCAST) definitive document E.DEF 7.3 Method (36) at Evotec (UK) Ltd (Manchester, UK). For testing by CLSI methodology, as per protocol, RPMI-1640 medium with 3-(*N*-morpholino) propanesulfonic acid (RPMI), an inoculum of 0.25~0.5 × 10^3^ CFU/well, and incubation at 35°C for 24h in ambient air was used. The results were read after 24-h incubation. MIC endpoints were determined visually as the lowest concentration of compound that resulted in a decrease of growth by ≥50% relative to that of the growth control (azole endpoint)(35). Stock solutions of test agents were prepared in neat DMSO and then diluted 100-fold to the desired concentrations with DMSO. The compound DMSO solution was applied to fungus growth media to ensure that the final concentration of DMSO was 1% (v/v) in all plates evaluated. For EUCAST, suspensions equivalent to a McFarland 0.5 standard were prepared for each test strain in 5 mL sterile water and diluted 1:10 in sterile water to provide an inoculum of 1-5 × 10^5^ CFU/mL. 100 μL of each inoculum was dispensed into all wells, except negative control wells (100 μL sterile water added), to provide 0.5-2.5 × 10^5^ CFU/well and a final volume of 200 μL single-strength RPMI-1640 was added per well. Assay plates were incubated according to EUCAST E.DEF 7.3 guidelines (35±2°C for 24±2 h in ambient air). An inoculum purity and viability check of the QC strains *C. krusei* ATCC 6258 and *C. parapsilosis* ATCC 22019 was performed on SABC agar. Compound preparation and treatment were conducted as shown above. The MIC was read as the lowest drug concentration that prevented any discernible growth (50% inhibition).

### Statistical analysis

For *in vitro* test data, statistical analyses of all the data were performed using the PRISM 8^®^ software program (GraphPad Software Inc., San Diego, CA, USA)., and the results are expressed as geo-metric mean (geo-mean), MIC_50_ (50 percentile) and MIC_90_ (90 percentile). Multiple comparison was performed by ANOVA followed by Turkey’s multiple comparison test. Statistical significance was defined as *p*<0.05.

For *in vivo* test data, survival analysis was performed by Kaplan-Meier plots followed by the log rank (Mantel-Cox) tests using the PRISM 8^®^ software program (GraphPad Software Inc., San Diego, CA, USA). All biomarker results are expressed as means ± standard error of the mean (SEM). CFU result was expressed as geometric means ± standard error (SD). Multiple comparison was performed by the Kruskal-Wallis with Dunn’s *post hoc* comparison using the PRISM 8^®^ software program (GraphPad Software Inc., San Diego, CA, USA). Comparison between 2 groups was performed by paired Wilcoxon paired rank test for MIC analysis or unpaired Mann-Whitney test *in vivo*. The ID_3-log_ value, which is the dose to reduce fungal load at 3-Log, was also calculated from dose-response curve using a nonlinear regression analysis with three parameter fitting using the PRISM 8^®^ software program. Statistical significance was defined as *P*<0.05.

## Results

### *Candida albicans* induced lung injury and death

Intranasal inoculation of Candida albicans (529L [ATCC^®^MYA4901™] strain) with either 2.5 × 10^6^/mouse or at 1.0 × 10^7^/mouse showed remarkable fungal loads in the lung and concentrations of biomarkers in BALF on Day 3 and 5 post inoculation (Fig 2). β-glucan in serum (as a marker of Candida infection) reached a peak on Day 5 at the lower inoculum, 2 days after the peak of fungal burden in the lung (Fig 2B). CXCL1 in BALF was increased on Day 3, and TNFα in serum peaked 2 days later, which showed similar kinetics with serum β-glucan. Thus, there is a time-gap between lung infection and subsequent systemic disseminated *Candida* infection. Considering biomarker data viability and loss of animals from the study, we decided to use 2.5 x10^6^/mouse inoculum and observe for 5 days, so that both survival and biomarkers could be analysed for pharmacological evaluation of PC945.

**Fig 2.**
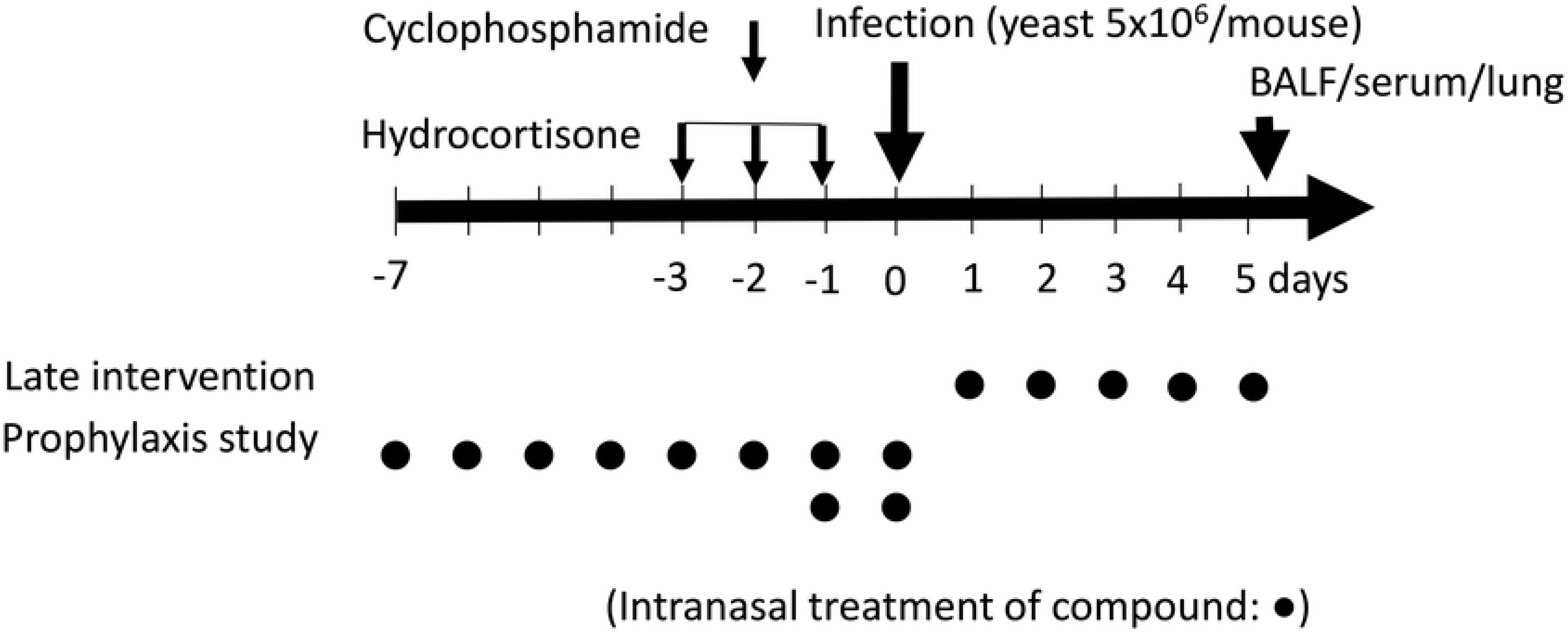
Time-dependent changes of fungal load in the lung (A), β-glucan in serum (B), CXCL-1 in BALF (C) and TNFα in serum (D) in A/J mice inoculated intranasally with (529L) at 2.5 x10^6^ yeasts/mouse or at 1.0 x10^7^ yeasts/mouse. CTL: non-infected control.

### Therapeutic antifungal effects of intranasal PC945 on *Candida albicans* infected mice

Prior to *in vivo* study, we evaluated the anti-fungal activities of PC945 and voriconazole against was *C. albicans* 529L, which was used for *in vivo* assay. In the broth microdilution assay [EUCAST method], PC945 showed an MIC value of 0.016 μg/mL, whereas the MIC for voriconazole was 0.008 μg/mL, suggesting that the *in vitro* effect of voriconazole was similar to that of PC945 against this strain.

As shown in Fig 1, PC945 was given intranasally daily a day post intranasal *Candida albicans* inoculation in immunocompromised, temporarily neutropenic mice. In this model, 71% of control mice (10/14) died or had dropped out of the study by day 5 post infection, so a minority (29%) of mice survived (Table 1, Fig 3A). Intranasally dosed PC945 saline suspension showed a dose-dependent improvement in survival rate (21, 36 and 79% at doses of 0.56, 2.8 and 14 μg/mouse, respectively) where the effect at the top dose was statistically significant (Table 1, Fig 3A). However, although intranasally dosed voriconazole also improved survival rate dose-dependently (Table 1), significant improvement was seen only at the highest dose (350μg/mouse), 25-fold higher than the dose of PC945, which achieved statistically significant improvement. When 50% survival improvement dose is calculated, PC945 (5.5 μg/mouse) was 6.6-fold more potent than voriconazole (36.3 μg/mouse) as shown in Fig 3B. Thus, PC945 was more effective than voriconazole in the *in vivo* system when dosed intranasally.

**Table 1.**
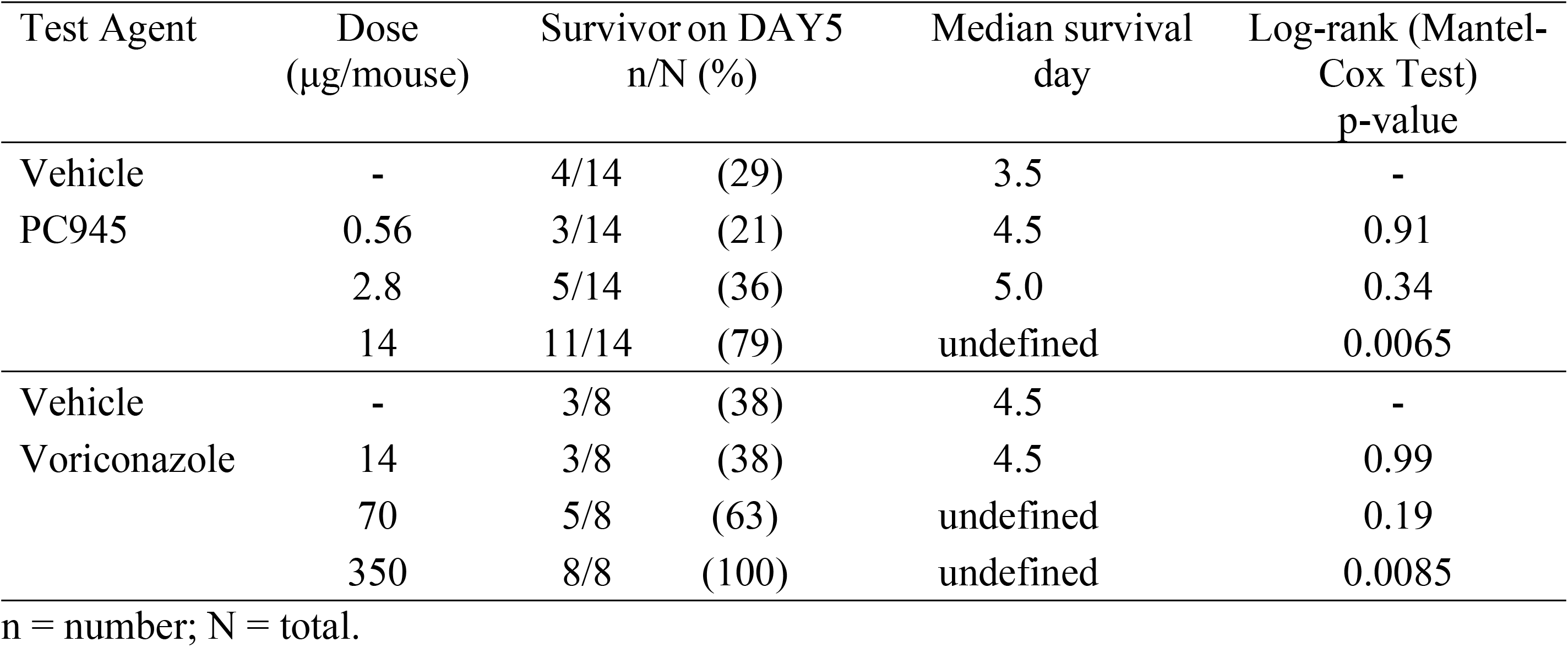
*In vivo* activities of effects of PC945 and voriconazole on in vivo survival on *C. albicans* infected mice

| Test Agent | Dose<br>($\mu\text{g}/\text{mouse}$) | Survivor on DAY5<br>n/N (%) | | Median survival<br>day | Log-rank (Mantel-Cox Test)<br>p-value |
| --- | --- | --- | --- | --- | --- |
| Vehicle | - | 4/14 | (29) | 3.5 | - |
| PC945 | 0.56 | 3/14 | (21) | 4.5 | 0.91 |
|  | 2.8 | 5/14 | (36) | 5.0 | 0.34 |
|  | 14 | 11/14 | (79) | undefined | 0.0065 |
| Vehicle | - | 3/8 | (38) | 4.5 | - |
| Voriconazole | 14 | 3/8 | (38) | 4.5 | 0.99 |
|  | 70 | 5/8 | (63) | undefined | 0.19 |
|  | 350 | 8/8 | (100) | undefined | 0.0085 |

n = number; N = total.

**Fig 3.**
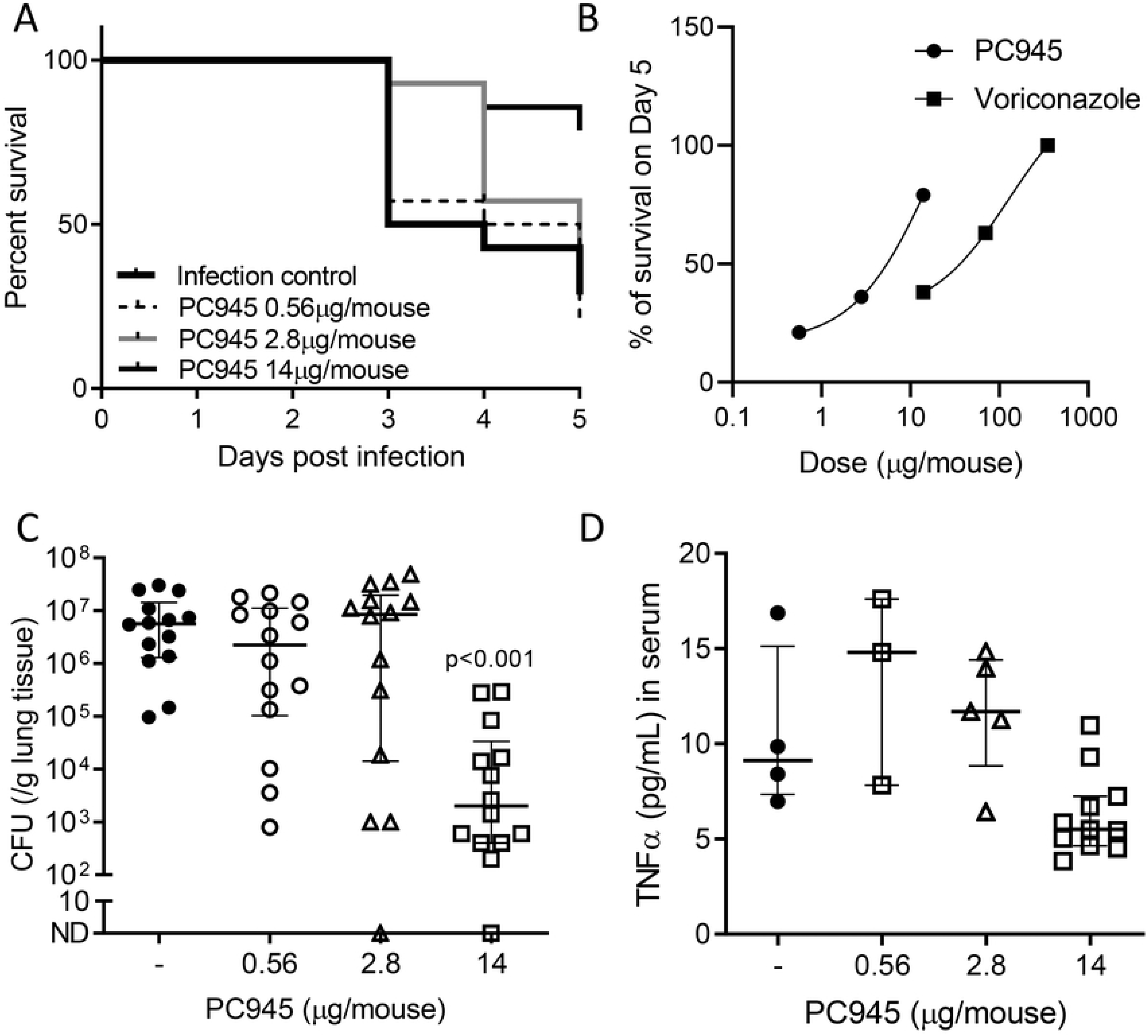
Antifungal activity of PC945 against *C. albicans in vivo*. Effect of once daily intranasal treatment of PC945 (0.56, 2.8, 14 μg/mouse) for 5 days on survival in *C. albicans* infected immunocompromised mice (N=14) (A) and a dose-response curve of survival rate for PC945 and voriconazole (14, 70, μg/mouse) (B). Lung fungal loads (CFU/g lung tissue) in all mice (C) and TNFα in serum in survived mice (D) were evaluated. Antifungal activity of extended prophylaxis treatment of PC945 was also evaluated. Each horizontal bar shows the geometric mean ± SD. * *p*<0.05.

In this model, high levels of fungal burden in lung (geometric mean: 6.6 ± 0.20 Log CFU/g of lung) were observed on day 5 in control mice, which PC945 dose-dependently inhibited as seen in Fig 3C. The dose which reduced fungal load by 3 Logs was 13.3 μg/mouse and 91.0 μg/mouse for PC945 and voriconazole, respectively, suggesting PC945 was 6.8-fold more potent than voriconazole. In addition, high levels of the inflammatory marker CXCL1 (5.3 ± 1.1 ng/mL) were detected in BALF on day 5 in control mice, which was significantly inhibited by PC945 at 14μg/mouse (Table 2). Serum TNFα was only evaluated in surviving mice, and PC945 showed a trend towards dose-dependent reduction (Table 2 and Fig 3D), which was not statistically significant due to the smaller sample size (Table 2).

**Table 2.**
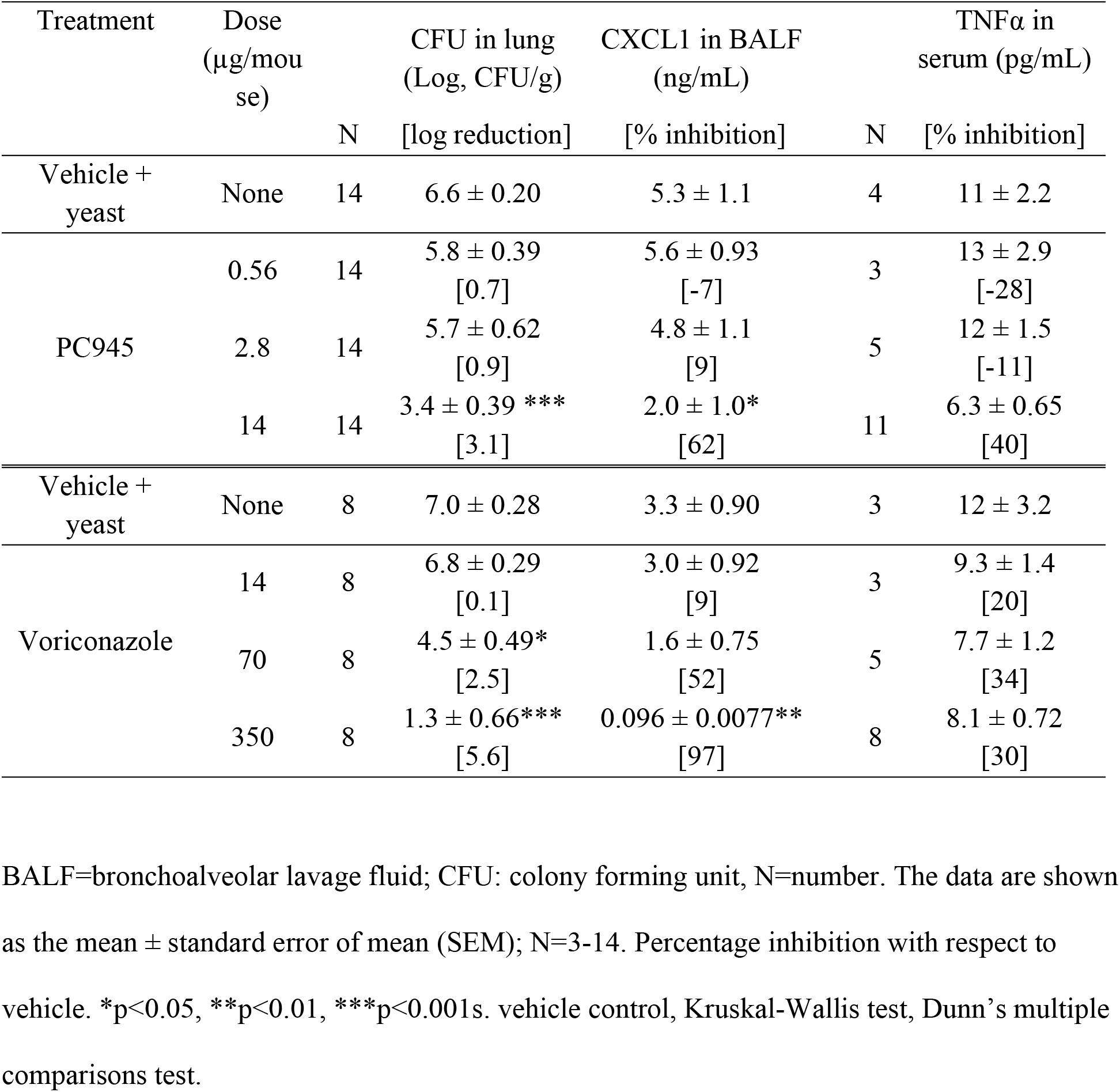
Effects of PC945 and voriconazole on biomarkers in BALF and serum obtained from *Candida albicans* infected, immuno-compromised, neutropenic mice

Histology revealed significant inflammation and minor acute lung injury as well as candida colonization (Grocott staining) on day 5 post infection or earlier (mortality) in control mice. PC945 completely eliminated candida from lung at 14 μg/mouse, except for animals where there was early death (Fig 4).

**Fig 4.**
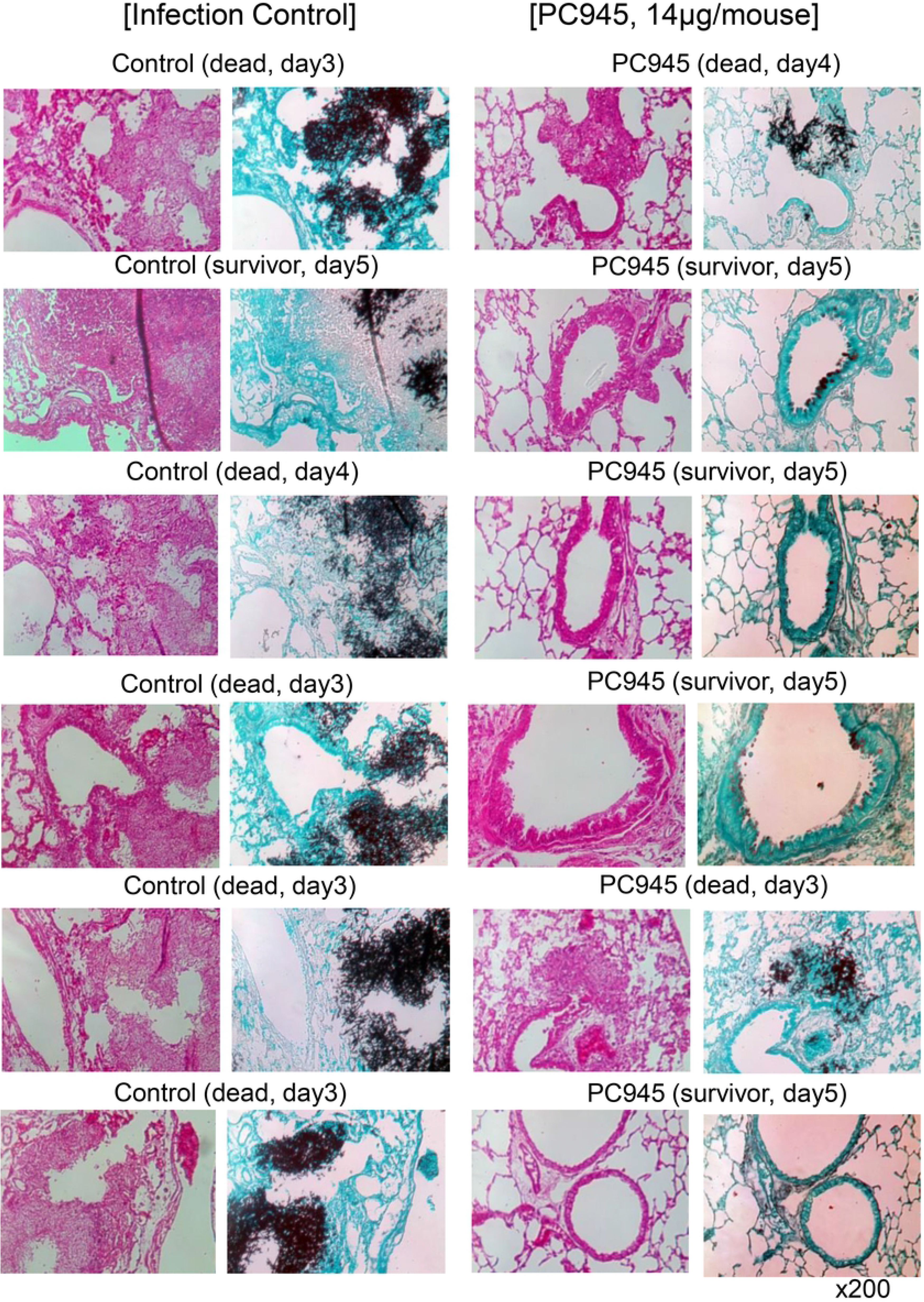
Histology of lung obtained from control and PC945-treated *C. albicans* infected immunocompromised mice. PC945 (14 μg/mouse) or saline was treated intranasally once daily, and some mice were dead before Day 5 post *C. albicans* inoculation and others were sacrificed on Day 5, 6 hrs after the last dose. Lung specimens were stained with Haematoxylin eosin (H&E) and Grocott fungus staining agent (x200).

### Antifungal effects of extended prophylaxis with intranasal PC945 on *Candida albicans* infected mice

As well as therapeutic dosing, the anti-fungal potential of PC945 using extended prophylaxis was evaluated (Fig 1). Extended prophylaxis (Day −7 to Day 0) at 0.56 μg/mouse showed marked and significant inhibition of fungal load in lung tissue (CFU) compared with shorter prophylaxis (Day −1 to Day 0) (Fig 5A). These effects were equivalent to those observed by therapeutic treatment of PC945 at 14 μg/mouse (Fig 3), suggesting extended prophylaxis was 25-fold more effective than therapeutic treatment. The extended prophylactic treatment also showed significant reduction of serum TNFα (Fig 5B).

**Fig 5.**
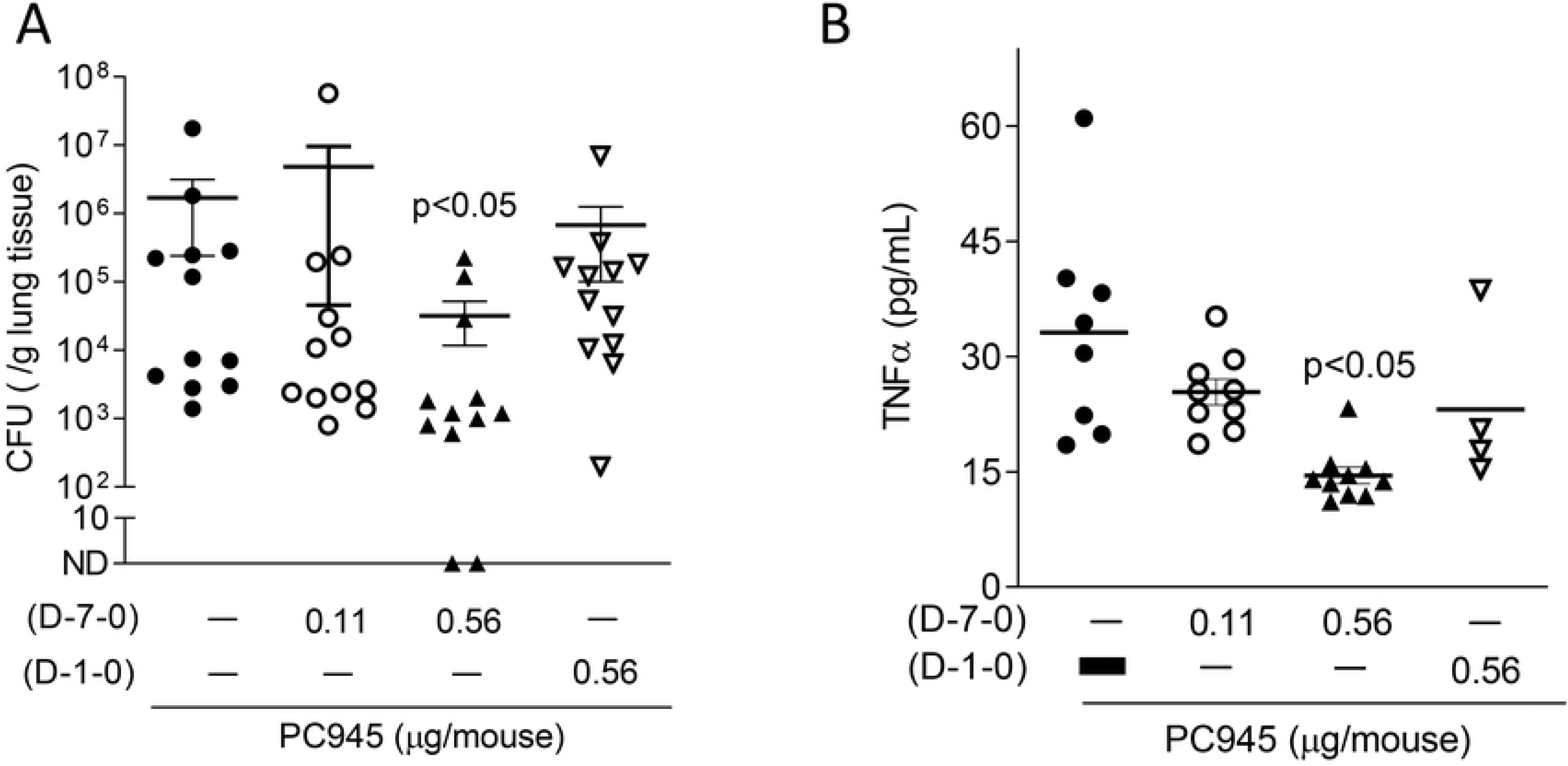
Antifungal activity of extended prophylaxis with PC945 against *C. albicans in vivo*. Effects of 8 days extended prophylaxis with intranasal PC945 (0.11, 0.56 μg/mouse, once daily) was compared with that of 2 days prophylaxis treatment only on lung fungal load (CFU/g tissue) (A) and TNFα in serum (B). Each horizontal bar shows the geometric mean ± SD. * *P*<0.05.

### *In vitro* antifungal activity against *Candida species*

The geometric mean MIC, MIC_50_ and MIC_90_ of PC945 against all *Candida* species tested by CLSI method were 0.027, 0.031 and 0.25 μg/mL, respectively, >2-fold weaker than voriconazole (≤0.016, ≤0.016, 0.063) (Table 3) and at least 2-fold more potent than posaconazole (0.097, 0.063, 0.5) (S2 Table). When analysing each *Candida* species separately, geometric mean MIC values of PC945 against *C. albicans* [37 isolates]*, C. tropicalis* [7]*, C. parapsilosis* [17]*, C. glabrata* [10], *C. lucitaniae* [11] and *C. guilliermondii* [6] were 0.017, 0.063, 0.017, 0.12, ≤0.016 and 0.18 μg/mL. PC945 was therefore equal to or less potent than voriconazole and more potent than posaconazole (Table 3, Fig 6A). All azoles used showed high MIC values for *C. tropicalis* FA1572 strain, which is known to be fluconazole resistant (Fig 6B). MIC values of voriconazole and posaconazole for the quality control strains [*C. krusei* (ATCC 6258), *C. parapsilosis* (ATCC 22019)] were within the expected MIC range (0.063-0.125 and 0.125-0.25 for *C. krusei*, 0.016 and 0.063-0.125 for *C. parapsilosis*, respectively).

**Table 3.**
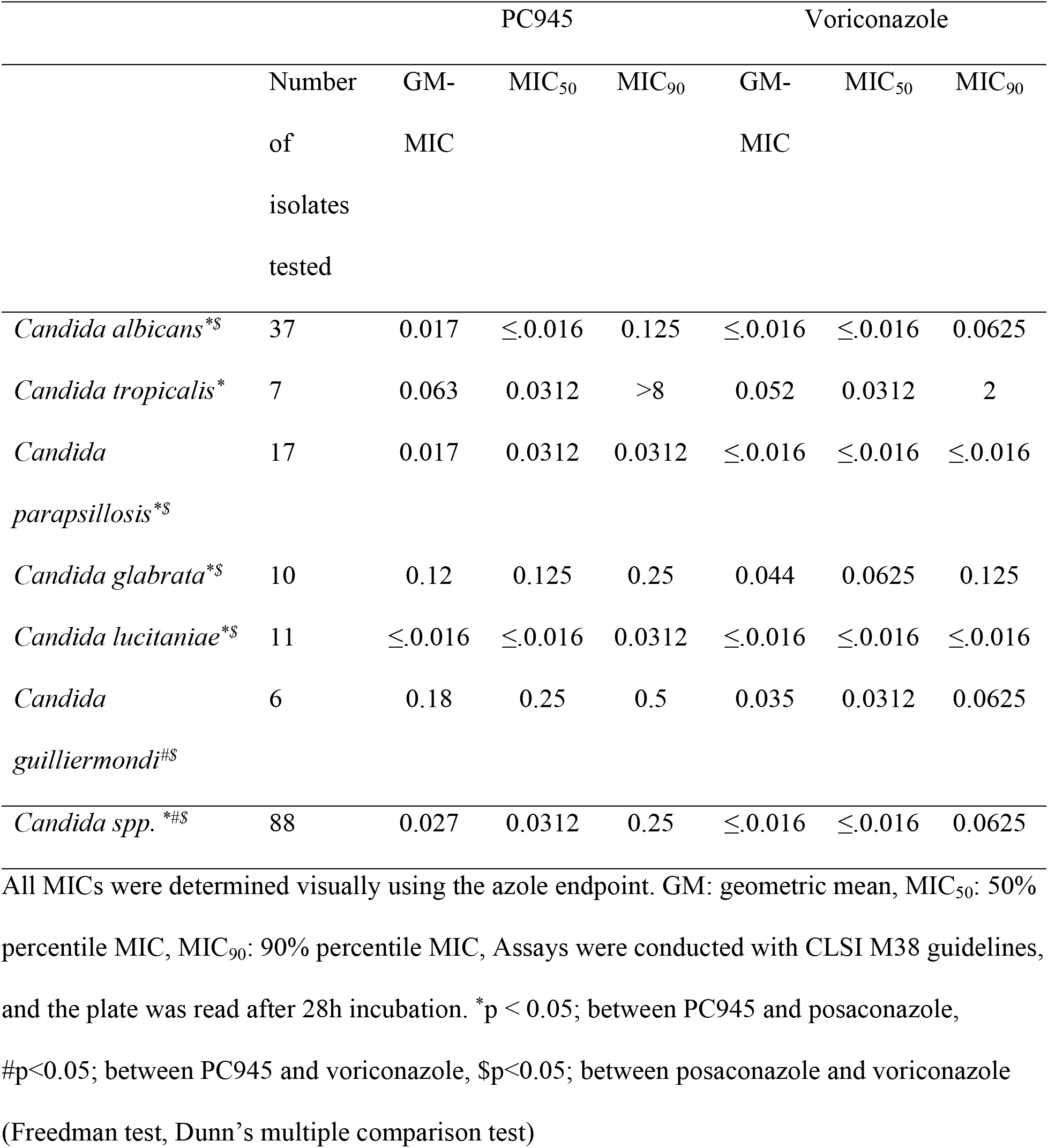
Susceptibility testing of *Candida spp.* to PC945 by CLSI

|  |  | PC945 |  |  | Voriconazole |  |  |
| --- | --- | --- | --- | --- | --- | --- | --- |
|  | Number<br>of<br>isolates<br>tested | GM-MIC | MIC<sub>50</sub> | MIC<sub>90</sub> | GM-MIC | MIC<sub>50</sub> | MIC<sub>90</sub> |
| *Candida albicans*<sup>*\$</sup> | 37 | 0.017 | ≤0.016 | 0.125 | ≤0.016 | ≤0.016 | 0.0625 |
| *Candida tropicalis*<sup>*</sup> | 7 | 0.063 | 0.0312 | >8 | 0.052 | 0.0312 | 2 |
| *Candida parapsilosis*<sup>*\$</sup> | 17 | 0.017 | 0.0312 | 0.0312 | ≤0.016 | ≤0.016 | ≤0.016 |
| *Candida glabrata*<sup>*\$</sup> | 10 | 0.12 | 0.125 | 0.25 | 0.044 | 0.0625 | 0.125 |
| *Candida lucitaniae*<sup>*\$</sup> | 11 | ≤0.016 | ≤0.016 | 0.0312 | ≤0.016 | ≤0.016 | ≤0.016 |
| *Candida guilliermondii*<sup>#\$</sup> | 6 | 0.18 | 0.25 | 0.5 | 0.035 | 0.0312 | 0.0625 |
| *Candida spp.*<sup>*\$</sup> | 88 | 0.027 | 0.0312 | 0.25 | ≤0.016 | ≤0.016 | 0.0625 |

All MICs were determined visually using the azole endpoint. GM: geometric mean, MIC<sub>50</sub>: 50% percentile MIC, MIC<sub>90</sub>: 90% percentile MIC, Assays were conducted with CLSI M38 guidelines, and the plate was read after 28h incubation. \*p < 0.05; between PC945 and posaconazole, #p<0.05; between PC945 and voriconazole, \$p<0.05; between posaconazole and voriconazole (Freedman test, Dunn's multiple comparison test)

**Fig 6.**
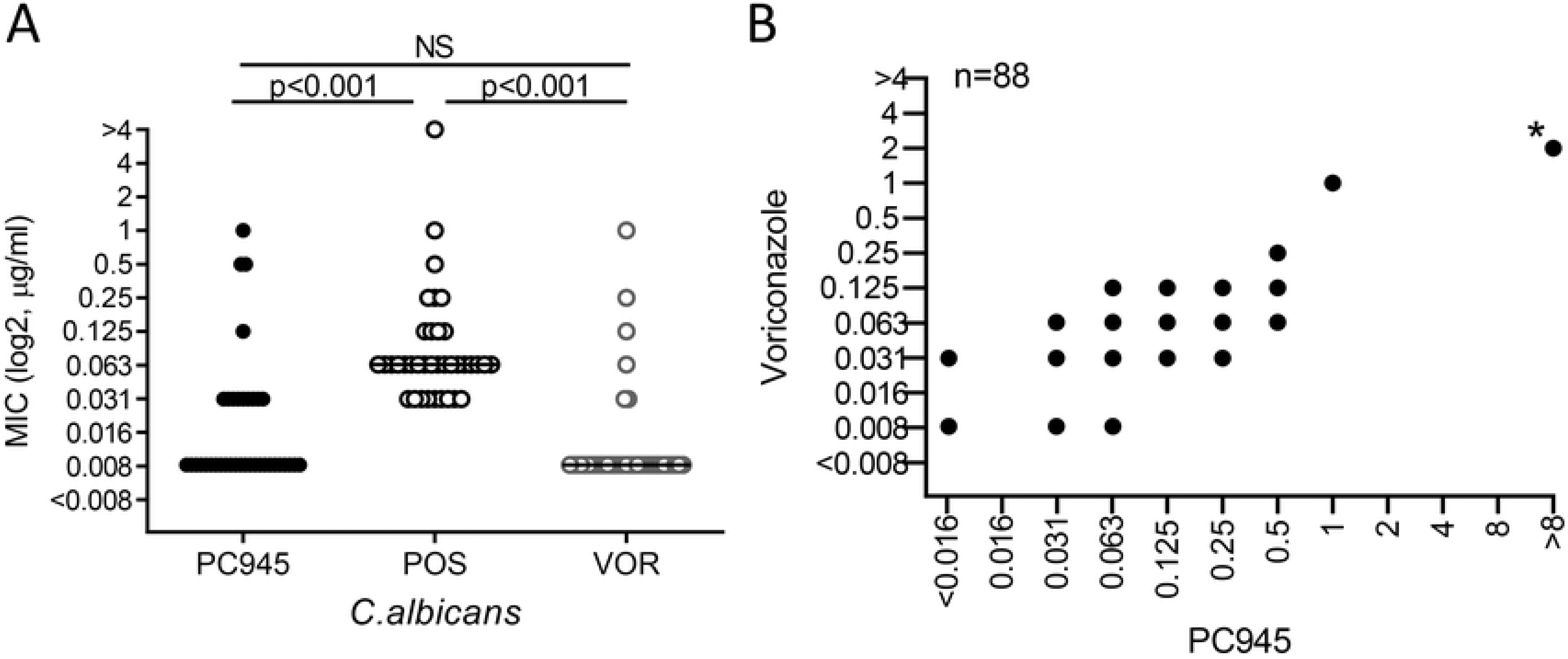
MIC distribution analyses: Individual MIC distribution versus *C. albicans* isolates (A) and the relationship of MIC distribution against all candida species between PC945 and Voriconazole (B) \**C. tropicalis* FA1572 strain

In addition, activities against same *Candida* isolates tested above were also evaluated using the EUCAST method. The geometric mean MIC, MIC_50_ and MIC_90_ values of PC945 against all *Candida* species tested were 0.045, 0.016 and 1 μg/mL, respectively, >2-fold less potent than voriconazole (S3 Table) as was observed using the CLSI method. There was a good correlation between the values from the CLSI and EUCAST tests (Pearson r=0.84, p<0.0001). MIC values of voriconazole versus the quality control strains [*C. krusei* (ATCC 6258), *C. parapsilosis* (ATCC 22019)] were within the expected MIC range (0.25 and 0.016, respectively).

## Discussion

PC945 has been found to be a potent antifungal agent, possessing activity against a broad range of *Candida* species, more potent than posaconazole, but comparable to or weaker than voriconazole in *in vitro* broth assays. *In vitro* anti-fungal activity of PC945 against the *C. albicans* (ATCC MYA4901 [529L]), used for in vivo analysis, was also slightly weaker than or similar to that of voriconazole. However, despite the weaker or similar *in vitro* potency, PC945 was 7~25-fold more potent than voriconazole in *C. albicans* infected mice *in vivo* when dosed intranasally once daily.

Although there is growing evidence that colonization of the lung by *C. albicans* impacts the pathogenesis of chronic respiratory diseases, experience with animal models to assess *in vivo* efficacy of antifungal agents is limited. Xu *et al.* reported a model where *C. albicans* was dosed intratracheally, showed acute lung injury and airway inflammation in both intact and immunosuppressed BALB/c mice, but the infection was only lethal in immunosuppressed mice (37). In this report, we used C5 deficient A/J mice which were found to be very susceptible to *A. fumigatus* infection (38, 39), with histology revealing significant airway inflammation and moderate acute lung injury.

In this model, PC945 was dosed to animals, followed 24 h later by inoculation with *C. albicans.* To evaluate biomarkers, all surviving animals were sacrificed on Day 5, so survival was not monitored thereafter. 71% of vehicle treated *C. albicans* infected control mice were withdrawn from the study by Day 5. However, once-daily treatment with PC945 showed dose-dependent beneficial effects on survival post inoculation with a statistically significant effect at 14 μg/mouse. Voriconazole showed statistically significant beneficial effects on survival at 350 μg/mouse, suggesting PC945 is much more potent than voriconazole *in vivo*. For fungal burden analysis (the dose inducing 3 Log reduction), PC945 was found to be 6.8-fold more potent than voriconazole. Despite displaying similar antifungal activities to voriconazole in broth microdilution assays *in vitro*, these results clearly indicate that PC945 significantly outperforms voriconazole *in vivo*.

This superior profile probably arises from several factors, including pharmacokinetics, physicochemical properties and unique pharmacological characteristics. As reported earlier, PC945 has been demonstrated to have a longer duration of action in bronchial epithelial cells and hyphae than voriconazole (29). Moreover, the molecule is retained within the lung, resulting in low systemic exposure after oral and face mask nebulization in a phase 1 study (NCT02715570) (40) (in preparation). Although patients have been treated with voriconazole by nebulization successfully (27), it required frequent dosing at extremely high doses due to its short duration of action in the lung and rapid systemic exposure after nebulization (27, 28). Voriconazole’s short pharmacodynamic duration of action in the lung relative to PC945 probably contributed to the relative antifungal activities seen here, where a once daily regimen was used. In addition, unique physicochemical properties might contribute to the beneficial character of PC945. As shown in S4 Table, PC945 is not Lipinski-compliant, with a high molecular weight, topological polar surface area (TPSA) and Log P value, so its physio-chemical properties are markedly different from those of voriconazole. PC945 also demonstrates lower aqueous solubility than voriconazole. As discussed above, PC945 has been optimised for a long residence time in airway epithelial cells and hyphae, but the molecular mechanism(s) leading to long lung residency and long lung duration of action have not been fully elucidated. Further studies are required to identify the mechanisms which produce PC945’s superior *in vivo* performance.

In addition, we have demonstrated here that 8-day prophylactic treatment (using very low doses) produced much greater anti-*Candida* activity than prophylactic treatment limited to 2 days. Furthermore, the effects of 8-day prophylactic treatment were maintained when treatment ceased just before *Candida albicans* inoculation on day 0 (Fig 5). This is powerful pharmacodynamic evidence that the effects of PC945 accumulate on daily dosing in mice and are maintained when dosing ceases. In fact, both our phase 1 study with healthy volunteers and rat studies demonstrated that PC945 accumulated in lung after repeat daily dosing (NCT02715570) (40) (in preparation). Interestingly, *Tolman* and colleagues have demonstrated that prophylaxis with aerosolized aqueous intravenous formulation of voriconazole significantly improved survival and limited the extent of invasive disease with *Aspergillus fumigatus*, as assessed by histopathology, in an invasive pulmonary murine model (26). Recently, Baistrocchi and colleagues reported that posaconazole accumulated in granulocyte type cells leading to enhanced anti-fungal effects (by exposure of cellular posaconazole during phagocytosis) (41). We have also demonstrated PC945 accumulation into alveolar cells in mice, including macrophages, after intranasal treatment (42) and showed that PC945 changes the cell integrity of *A. fumigatus* to accelerate elimination by macrophages/neutrophils (43). It is possible that loading of PC945 into granulocyte/macrophages might enhance the compound’s intrinsic anti-fungal activity against *candida albicans*.

Against 88 *Candida* isolates, there was a good correlation between the MIC values of PC945 and those of voriconazole. One isolate, *C. tropicalis* FA1572, which is known to be fluconazole resistant with a mutation of CYP51 at Y132 (44, 45), was found to be resistant to both treatments. Between species, the activities of PC945 against *C. albicans*, *C. tropicalis, C. parapsilosis* and *C. lucitaniae* were more potent than those against *C. glabrata* and *C. guilliermondi*. As previously reported, PC945 also inhibited the growth of *Candida auris* (n=72) (32) and *C. krusei* (n=1) (33). The effects of PC945, determined by the CLSI method, were confirmed using the EUCAST method. MICs against Candida species obtained by the CLSI and EUCAST methods are reported to show a good correlation for fluconazole and voriconazole in general, whereas MICs obtained by the EUCAST method are typically lower for posaconazole (47) than those obtained using CLSI. For PC945, there was a good correlation between results obtained using the two methods.

This study had several limitations. Firstly, this model is unlike natural chronic respiratory colonization seen in asthma or COPD, more closely resembling an immunocompromised setting (invasive form). We did not evaluate clinically relevant markers, such as lung function or symptoms, as readouts. Therefore, it may be inappropriate to extrapolate the results of this study directly to chronic respiratory disease with persistent Candida colonization in the clinic. However, we believe that this model is still useful, allowing assessment of the anti-fungal, pharmacodynamic properties of PC945. Secondly, there was a lack of pharmacokinetic measurements of PC945 in mice used in the current experiments to help understand the superior anti-fungal effects of PC945 versus voriconazole. Preliminary data suggested accumulation of PC945 in the lung after repeat intranasal treatment in *A. fumigatus* infected mice (42). As we used mice in the same condition (immunocompromised, temporally neutropenic) as this previous study, we expect similar pharmacokinetics in the current experiment. Thirdly, we showed CFU data as per gram of lung tissue. There might be many confounders that influence organ weight, such as condition of mice (oedema etc), variable volume of blood remained etc. However, we were not able to normalize to dried tissue weight as we had to get yeast from fresh lung tissue. As the difference of CFU between groups is >3 log, those factors are only likely to have a limited impact on the results.

Thus, we showed superior *in vivo* performance of PC945 versus voriconazole on fungal load and survival rate after intranasal treatment against *C. albicans* lung infection, replicating our findings using *Aspergillus fumigatus* infection (38). Further data suggested that the antifungal effects in the lung of PC945 accumulated on repeat dosing and that these effects were persistent. Thus, PC945 has the potential to be a novel inhaled therapy for the prevention or treatment of *C. albicans* lung infection in humans. PC945 is in clinical development (40, 48), and these *in vivo* data indicate that further clinical evaluation is warranted.

## ACKNOWLEDGEMENTS

We are grateful to Mr. Takahiro Nakaoki (Nihon Univ.) for assistance in the *in vivo* study. We also thank Dr. Mihiro Sunose, Syganture Discovery Ltd. (Nottingham, UK) for her assistance in predicting the water solubility of PC945. This work was financially supported by Pulmocide Ltd.

## Supporting information

**S1 Table. Dose unit conversion**

**S2 Table. Susceptibility testing of***Candida spp.* **to posaconazole by CLSI**

**S3 Table. Susceptibility testing of***Candida spp.* **to PC945, voriconazole by EUCAST**

**S4 Table. Physicochemical properties of PC945 and voriconazole**

## REFERENCE

1. Pappas PG, Kauffman CA, Andes D, Benjamin DK, Jr., Calandra TF, Edwards JE, Jr., et al. Clinical practice guidelines for the management of candidiasis: 2009 update by the Infectious Diseases Society of America. Clin Infect Dis. 2009;48(5):503–35.

2. Johnson DC. Chronic candidal bronchitis: a consecutive series. Open Respir Med J. 2012;6:145–9.

3. Maiz L, Nieto R, Canton R, Gomez GdlPE, Martinez-Garcia MA. Fungi in Bronchiectasis: A Concise Review. Int J Mol Sci. 2018;19(1).

4. Nguyen LD, Viscogliosi E, Delhaes L. The lung mycobiome: an emerging field of the human respiratory microbiome. Front Microbiol. 2015;6:89.

5. Yazici O, Cortuk M, Casim H, Cetinkaya E, Mert A, Benli AR. Candida glabrata Pneumonia in a Patient with Chronic Obstructive Pulmonary Disease. Case Rep Infect Dis. 2016;2016:4737321.

6. Dermawan JKT, Ghosh S, Keating MK, Gopalakrishna KV, Mukhopadhyay S. Candida pneumonia with severe clinical course, recovery with antifungal therapy and unusual pathologic findings: A case report. Medicine (Baltimore). 2018;97(2):e9650.

7. Ramachandran S, Shah A, Pant K, Bhagat R, Jaggi OP. Allergic bronchopulmonary aspergillosis and Candida albicans colonization of the respiratory tract in corticosteroid-dependent asthma. Asian Pac J Allergy Immunol. 1990;8(2):123–6.

8. Mullaoglu S, Turktas H, Kokturk N, Tuncer C, Kalkanci A, Kustimur S. Esophageal candidiasis and Candida colonization in asthma patients on inhaled steroids. Allergy Asthma Proc. 2007;28(5):544–9.

9. Chowdhary A, Agarwal K, Kathuria S, Gaur SN, Randhawa HS, Meis JF. Allergic bronchopulmonary mycosis due to fungi other than Aspergillus: a global overview. Crit Rev Microbiol. 2014;40(1):30–48.

10. O’Driscoll BR, Hopkinson LC, Denning DW. Mold sensitization is common amongst patients with severe asthma requiring multiple hospital admissions. BMC Pulm Med. 2005;5:4.

11. Maiz L, Vendrell M, Olveira C, Giron R, Nieto R, Martinez-Garcia MA. Prevalence and factors associated with isolation of Aspergillus and Candida from sputum in patients with non-cystic fibrosis bronchiectasis. Respiration. 2015;89(5):396–403.

12. Barchiesi F, Orsetti E, Gesuita R, Skrami E, Manso E, Candidemia Study G. Epidemiology, clinical characteristics, and outcome of candidemia in a tertiary referral center in Italy from 2010 to 2014. Infection. 2016;44(2):205–13.

13. van der Geest PJ, Dieters EI, Rijnders B, Groeneveld JA. Safety and efficacy of amphotericin-B deoxycholate inhalation in critically ill patients with respiratory Candida spp. colonization: a retrospective analysis. BMC infectious diseases. 2014;14:575.

14. Cottier F, Hall RA. Face/Off: The Interchangeable Side of Candida Albicans. Front Cell Infect Microbiol. 2019;9:471.

15. Morales DK, Hogan DA. Candida albicans interactions with bacteria in the context of human health and disease. PLoS Pathog. 2010;6(4):e1000886.

16. Roux D, Gaudry S, Dreyfuss D, El-Benna J, de Prost N, Denamur E, et al. Candida albicans impairs macrophage function and facilitates Pseudomonas aeruginosa pneumonia in rat. Crit Care Med. 2009;37(3):1062–7.

17. Roux D, Gaudry S, Khoy-Ear L, Aloulou M, Phillips-Houlbracq M, Bex J, et al. Airway fungal colonization compromises the immune system allowing bacterial pneumonia to prevail. Crit Care Med. 2013;41(9):e191–9.

18. De Pascale G, Antonelli M. Candida colonization of respiratory tract: to treat or not to treat, will we ever get an answer? Intensive Care Med. 2014;40(9):1381–4.

19. Thompson GR, 3rd, Lewis JS, 2nd. Pharmacology and clinical use of voriconazole. Expert Opin Drug Metab Toxicol. 2010;6(1):83–94.

20. Xiong WH, Brown RL, Reed B, Burke NS, Duvoisin RM, Morgans CW. Voriconazole, an antifungal triazol that causes visual side effects, is an inhibitor of TRPM1 and TRPM3 channels. Invest Ophthalmol Vis Sci. 2015;56(2):1367–73.

21. Jeong S, Nguyen PD, Desta Z. Comprehensive in vitro analysis of voriconazole inhibition of eight cytochrome P450 (CYP) enzymes: major effect on CYPs 2B6, 2C9, 2C19, and 3A. Antimicrob Agents Chemother. 2009;53(2):541–51.

22. Bruggemann RJ, Donnelly JP, Aarnoutse RE, Warris A, Blijlevens NM, Mouton JW, et al. Therapeutic drug monitoring of voriconazole. Ther Drug Monit. 2008;30(4):403–11.

23. Rodvold KA, Yoo L, George JM. Penetration of anti-infective agents into pulmonary epithelial lining fluid: focus on antifungal, antitubercular and miscellaneous anti-infective agents. Clin Pharmacokinet. 2011;50(11):689–704.

24. Drusano GL. Antimicrobial pharmacodynamics: critical interactions of ‘bug and drug’. Nature reviews Microbiology. 2004;2(4):289–300.

25. Capitano B, Potoski BA, Husain S, Zhang S, Paterson DL, Studer SM, et al. Intrapulmonary penetration of voriconazole in patients receiving an oral prophylactic regimen. Antimicrob Agents Chemother. 2006;50(5):1878–80.

26. Tolman JA, Wiederhold NP, McConville JT, Najvar LK, Bocanegra R, Peters JI, et al. Inhaled voriconazole for prevention of invasive pulmonary aspergillosis. Antimicrob Agents Chemother. 2009;53(6):2613–5.

27. Hilberg O, Andersen CU, Henning O, Lundby T, Mortensen J, Bendstrup E. Remarkably efficient inhaled antifungal monotherapy for invasive pulmonary aspergillosis. Eur Respir J. 2012;40(1):271–3.

28. Andersen CU, Sonderskov LD, Bendstrup E, Voldby N, Cass L, Ayrton J, et al. Voriconazole Concentrations in Plasma and Epithelial Lining Fluid after Inhalation and Oral Treatment. Basic & clinical pharmacology & toxicology. 2017;121(5):430–4.

29. Colley T, Alanio A, Kelly SL, Sehra G, Kizawa Y, Warrilow AG, et al. In vitro and in vivo antifungal profile of a novel and long acting inhaled azole, PC945, on Aspergillus fumigatus infection. Antimicrob Agents Chemother. 2017.

30. Colley T, Ito K, Rapeport G, Strong P, Murray PJ, Onions S, et al. Antimycotic compound. 2016:WO2016087880.

31. Pagani N, Murray, A., Strong, P., Ito, K., Rapeport, G., Cass, L., Carby, M., Simon, A., Armstrong-James, D., Reed, A. Use of a novel inhaled azole for treatment for fungal tracheobronchitis post-lung transplantation: a case report. Journal of Fungi. 2019(Fungal Update, abstract,):in press.

32. Rudramurthy SM, Colley T, Abdolrasouli A, Ashman J, Dhaliwal M, Kaur H, et al. In vitro antifungal activity of a novel topical triazole PC945 against emerging yeast Candida auris. The Journal of antimicrobial chemotherapy. 2019;74(10):2943–9.

33. Colley T, Alanio A, Kelly SL, Sehra G, Kizawa Y, Warrilow AGS, et al. In Vitro and In Vivo Antifungal Profile of a Novel and Long-Acting Inhaled Azole, PC945, on Aspergillus fumigatus Infection. Antimicrob Agents Chemother. 2017;61(5):e02280–16.

34. Kimura G, Ueda K, Eto S, Watanabe Y, Masuko T, Kusama T, et al. Toll-like receptor 3 stimulation causes corticosteroid-refractory airway neutrophilia and hyperresponsiveness in mice. Chest. 2013;144(1):99–105.

35. Institute CaLS. Reference method for broth dilution antifungal susceptibility testing of yeasts, 3rd ed Approved standard M27-A3. Clinical and Laboratory Standards Institute, Wayne, PA; 2008.

36. Arendrup MC, Guinea, J., Cuenca-Estrella, M., Meletiadis, J., Mouton, J.W., Lagrou, K., Howard, S.J., the Subcommittee on Antifungal Susceptibility Testing (AFST) of the ESCMID European Committee for, (EUCAST) AST. EUCAST DEFINITIVE DOCUMENT E.DEF 7.3, Method for the determination of broth dilution minimum Inhibitory concentrations of antifungal agents for yeasts. EUCAST-AFST; 2015.

37. Xu ZL, Li SR, Fu L, Zheng L, Ye J, Li JB. Candida albicans-induced acute lung injury through activating several inflammatory signaling pathways in mice. Int Immunopharmacol. 2019;72:275–83.

38. Kimura G, Nakaoki T, Colley T, Rapeport G, Strong P, Ito K, et al. In Vivo Biomarker Analysis of the Effects of Intranasally Dosed PC945, a Novel Antifungal Triazole, on Aspergillus fumigatus Infection in Immunocompromised Mice. Antimicrob Agents Chemother. 2017;61(9):e00124–17.

39. Zaas AK, Liao G, Chien JW, Weinberg C, Shore D, Giles SS, et al. Plasminogen Alleles Influence Susceptibility to Invasive Aspergillosis. PLoS Genet. 2008;4(6):e1000101.

40. ClinicalTiralsgov. A Study to Investigate the Safety, Tolerability and Pharmacokinetics of Single and Repeat Doses of PC945, NCT02715570 2018 [Available from: https://clinicaltrials.gov/ct2/show/NCT02715570?term=PC945&rank=1.

41. Baistrocchi SR, Lee MJ, Lehoux M, Ralph B, Snarr BD, Robitaille R, et al. Posaconazole-Loaded Leukocytes as a Novel Treatment Strategy Targeting Invasive Pulmonary Aspergillosis. J Infect Dis. 2016.

42. Ito K, Kizawa, Y., Kimura, G., Nishimoto, Y., Rapeport, G., Strong, P. Accumulation of a novel inhaled azole, PC945 in alveolar cells in temporally neutropenic immunocompromised mice infected with Aspergillus fumigatus AAAM2020; Lugano2020.

43. Armstrong-James D, Colley, T., Strong, P., Rapeport, G., Ito, K. Altered A.fumigatus cell wall integrity by PC945, a novel inhaled azole AAAM2020; Lugano2020.

44. Warn PA, Morrissey J, Moore CB, Denning DW. In vivo activity of amphotericin B lipid complex in immunocompromised mice against fluconazole-resistant or fluconazole-susceptible Candida tropicalis. Antimicrob Agents Chemother. 2000;44(10):2664–71.

45. Castanheira M, Deshpande LM, Messer SA, Rhomberg PR, Pfaller MA. Analysis of global antifungal surveillance results reveals predominance of Erg11 Y132F alteration among azole-resistant Candida parapsilosis and Candida tropicalis and country-specific isolate dissemination. Int J Antimicrob Agents. 2020;55(1):105799.

46. Warn PA. ASSESSMENT OF THE IN VIVO EFFICACY OF CHD-FA (CarboHydrate Derived - Fulvic Acid) IN A MURINE MODEL OF DISSEMINATED CANDIDIASIS https://www.fulvicacid.co.za: Euprotec Ltd.; 2008.

47. Pfaller MA, Castanheira M, Messer SA, Rhomberg PR, Jones RN. Comparison of EUCAST and CLSI broth microdilution methods for the susceptibility testing of 10 systemically active antifungal agents when tested against Candida spp. Diagn Microbiol Infect Dis. 2014;79(2):198–204.

48. The Effect of PC945 on Aspergillus Fumigatus Lung Infection in Patients With Asthma, NCT03745196 [Internet]. 2018.

